# Spontaneous eye blinks as temporal markers of internal attention

**DOI:** 10.64898/2026.07.06.736774

**Authors:** Daniel Schneider, Şahcan Özdemir, Edmund Wascher, Stefan Arnau

## Abstract

While eye blinks interrupt the flow of visual information, their timing may reveal the temporal organisation of cognitive processing. Here, we asked whether the temporal distribution of blinks across trials provides a time-resolved signature of attentional focusing in working memory. In Experiment 1, blink-locked EEG analyses showed that blink timing was aligned with neural activity reflecting attentional focusing on a relevant memory representation. In Experiment 2, participants remembered the same visual information across conditions, but the relevant item was cued either early during storage, or later with presentation of the memory probe. Blink-frequency profiles shifted accordingly, increasing after the cue when selection was possible early and after the probe when selection was delayed. Post-cue blinks in the early-selection condition were also associated with better memory performance. Thus, our findings establish eye-blink frequency profiles as an unobtrusive chronometric signal for tracking latent cognitive processing in settings where neuroimaging is not feasible.

## Main

Traditionally, spontaneous blinks have been conceptualized primarily as peripheral maintenance behaviours serving ocular lubrication and protection, with their occurrence assumed to be largely random or governed by low-level physiological rhythms. However, this view is difficult to reconcile with evidence showing that blink timing is systematically modulated by cognitive processing demands.^1^ Rather than merely interrupting visual input, spontaneous blinks appear to occur at specific moments within ongoing cognition, suggesting that their temporal distribution may provide information about when cognitive operations are initiated, completed, or reorganized.

Research across a wide range of paradigms demonstrates that spontaneous blinks exhibit a non-random temporal structure. Blink rate is reduced during sustained visual attention, and blinks are suppressed while task-relevant information is expected or actively processed.^2,3^ Complementing this, blink latencies have been shown to vary with the temporal window during which visual information is relevant for behaviour, suggesting that blink timing is closely coordinated with periods of external information sampling.^4,5^ This temporal coordination has been particularly well documented in reading and visual word recognition. During natural reading, blinks tend to cluster around punctuation marks, sentence boundaries, and semantic breakpoints, indicating that they occur preferentially at moments of local processing completion or transition between units of comprehension.^5–8^ Similar patterns have been observed during continuous speech perception and film viewing ^9^, where blinks align with narrative boundaries or low-information segments.^10^ Related evidence comes from an anti-saccade task, in which time-resolved blink-probability varied with trial type and performance, suggesting sensitivity not only to continuous sampling of sensory input but to upcoming inhibitory control demands.^11^

Together, these findings suggest that blink timing is aligned with transitions in ongoing processing, with blinks occurring when external information sampling can be briefly reduced and being suppressed when task-relevant information is imminent. This temporal sensitivity may make the distribution of blinks particularly valuable for studying working memory, where transitions in representational states occur as internally maintained information is selected, prioritised, and transformed in preparation for subsequent cognitive operations or actions. Blink timing may therefore provide a chronometrically informative signal of when such latent operations occur, even after the relevant sensory input is no longer available.

Initial evidence suggests that spontaneous blinking is systematically related to distinct phases and operations of working memory. Rac-Lubashevsky and colleagues ^12^ used an event-based eye-blink rate measure and showed that blink frequency increased during phases related to working memory updating. Ortega and colleagues ^13^ further showed that spontaneous blink rate varies across phases of a visual working memory task and reported a positive association between blink rate during the delay period and task performance. These findings show that spontaneous blinking varies systematically with working memory demands. However, for the distribution of blink times across trials to be informative about the timing of working memory operations, it must be shown that blinks are temporally aligned, for example, with neural markers of those operations, rather than merely reflecting broader constructs like task engagement or mental effort.

Against this background, the present study addressed whether spontaneous blink timing is systematically related to latent cognitive processes on the level of working memory. In Experiment 1, a working memory task with a large number of trials served as an experimental framework for relating blink timing to established EEG markers of attentional focusing and active storage of visuo-spatial information. Experiment 2 tested whether blinks alone can serve as a behavioural index of when attentional focusing is implemented within a working memory task. We examined whether their temporal distribution could recover the timing of this process without relying on concurrent neural measures. Participants performed a working memory task in which the same sensory input had to be maintained, but relevant information could be selected either early, with the presentation of a retrospective cue (retro-cue) before report, or only later, at the moment of report. If blink-frequency profiles can be used to infer the timing of working memory operations, their temporal pattern should follow this experimental manipulation: blink frequency should increase after the retro-cue when the relevant representation can be selected before report, but shift to the probe when this selection is delayed. Such a shift would indicate that spontaneous blink timing is chronometrically informative about when attentional selection occurs within the working memory task structure.

More generally, demonstrating that blink timing reflects how attentional selection unfolds within working memory would support its value as a simple, unobtrusive, and easily measurable chronometric signal with potential relevance across cognitive domains. This approach could be particularly useful for studying cognition in contexts where more complex neural measurements such as electroencephalography (EEG) or magnetoencephalography (MEG) are impractical, unavailable, or liable to interfere with task execution.

## Results

### Experiment 1: Spontaneous blinks are temporally aligned to attentional focusing in working memory

In Experiment 1, 32 participants performed a visual working memory task in which two oriented stimuli were briefly presented to the left and right of fixation and had to be maintained for later report (see Fig. 1a). After an encoding interval, a retro-cue indicated which of the two remembered orientations was relevant for the first report. Participants then adjusted a centrally presented probe orientation to match the cued item. After a further attentional re-orienting interval, they reported the remaining orientation.

**Figure 1.**
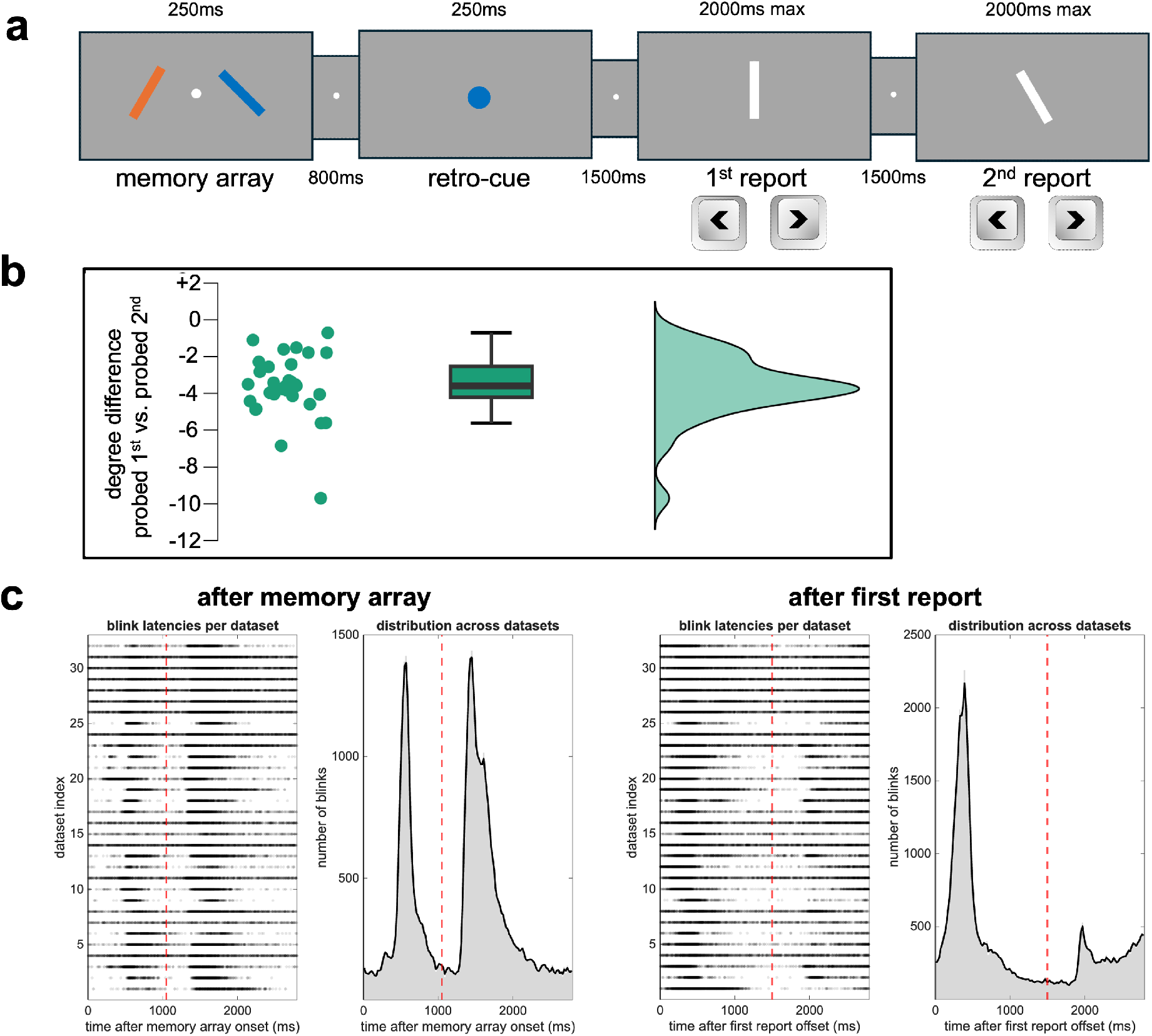
Design and blink latencies for Experiment 1. **(a)** The experimental design of Experiment 1, with a retro-cue providing information about the order of orientation reports. **(b)** The behavioural results (angular error / degree difference in orientation report), based on individual datapoints, a standard boxplot and a smoothed distribution plot of the first minus second item reports. **(c)** Temporal distribution of eye blinks for each dataset (one dot per blink) and pooled distributions across datasets (20 ms bins; smoothed curves are shown for visualisation). The left panels show the interval following memory array onset, with dashed red lines indicating retro-cue onset. The right panels show the interval following the report of the first item, with dashed red lines indicating the onset of the second memory probe.

Overall, participants performed the task reliably, with a lower angular error in reporting the relevant orientation for the first report position (M=11.66°, SE=0.364°, range = 8.0°-15.5°) than for the second position (M=15.24°, SE=0.492°, range = 15.1°-22.5°), t(31)=11.767, p<.001, dz=2.080, CI95 [2.959, 4.201]. This indicates a behavioural advantage for the cued orientation, which was also the orientation reported first (see Fig. 1b).

Spontaneous blinks occurred throughout the trial but were not uniformly distributed over time. Blink frequency was reduced around periods of external stimulation and report, and increased during extended intervals in which no new visual information had to be processed in working memory (see Fig. 1c). To determine whether blink timing was specifically aligned with working memory operations, we examined lateralised EEG activity time-locked to individual blinks.

The primary focus of the present analyses was on blink-locked lateralised EEG activity. Before conducting these analyses, however, we characterised the stimulus-locked temporal structure of lateralised EEG activity in the same set of trials (i.e., the trials containing the blinks subsequently included in the blink-locked analyses). We focused on two time windows: the first window after the retro-cue was critical because it was the period in which the cued item could be selected for the upcoming report. The second window encompassed the completion of the first orientation report and the subsequent attentional re-orienting towards the second item. These results show when the lateralised EEG effects occurred across the task and provide an important context for the subsequent blink-locked analyses.

All posterior lateralised EEG effects were analysed over electrode pairs PO7/8, P7/8, PO3/4, and P5/6, contrasting activity contralateral and ipsilateral to the location of the memory item relevant for the subsequent report. Depending on the analysed time window, lateralisation was therefore defined relative to the item required for the first or second orientation report.

Posterior alpha power lateralisation served as an index of attentional prioritisation of the relevant internal representation and appeared as a stronger suppression of oscillatory power contralateral to the location of the relevant working memory item.^14,15^ Following retro-cue presentation, this effect emerged with a maximum effect latency of around 500 ms (see Fig. 2a) and was followed by a CDA-like posterior ERP lateralisation (CDA: contralateral delay activity, see Fig. 2b). The CDA effect is a sustained contralateral negativity in the ERP that has been linked to the active storage of visuospatial information in working memory.^16^

**Figure 2.**
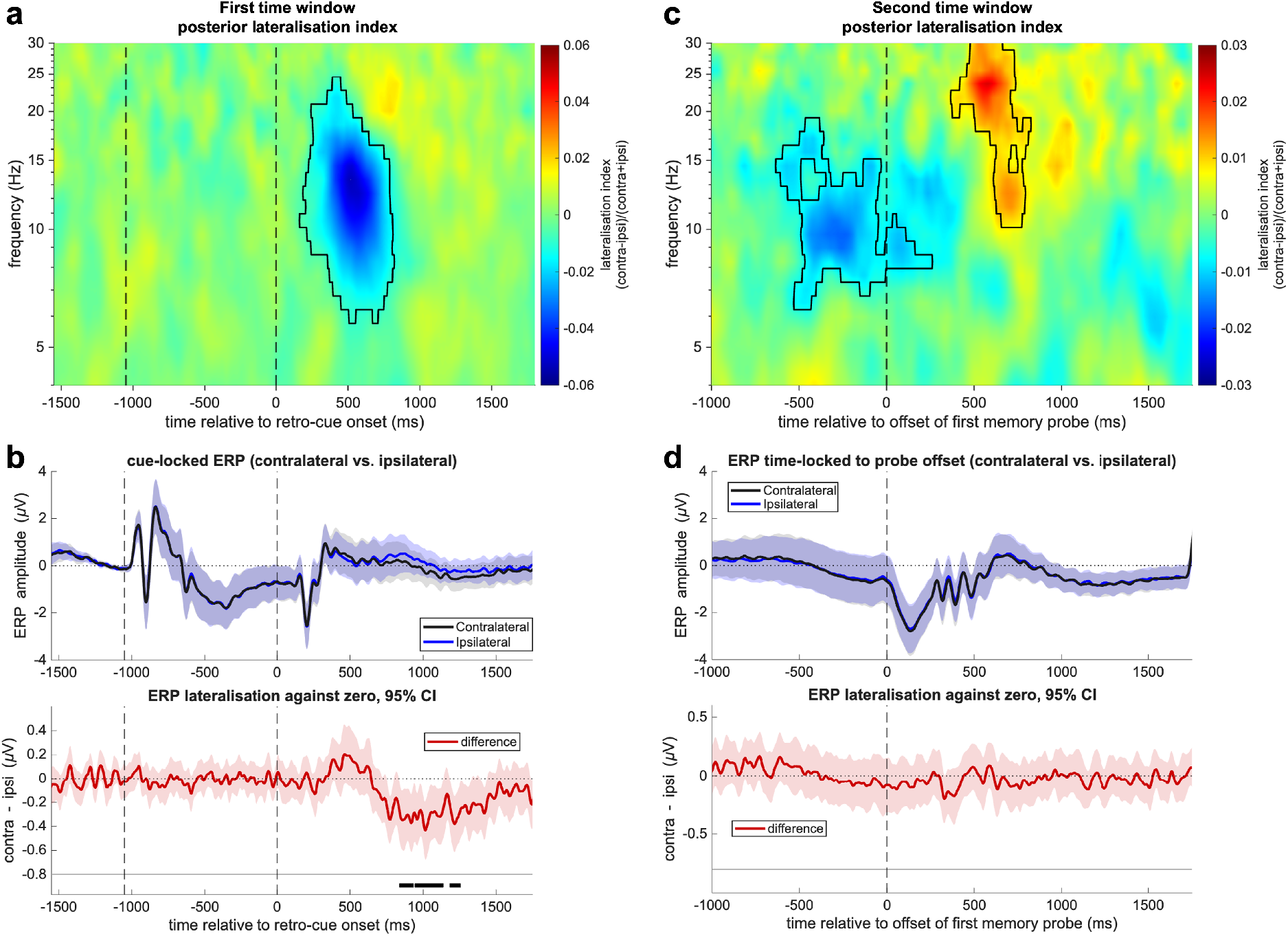
Stimulus-locked lateralised EEG effects in Experiment 1. Panels show stimulus-locked lateralised EEG activity in the two analysis windows used for the blink-locked analyses. **(a)** Posterior time-frequency lateralisation index relative to the side of the cued working memory item and time-locked to retro-cue onset. A pronounced posterior alpha lateralisation (contralateral suppression) emerged after the retro-cue. **(b)** Lateralised ERP activity time-locked to retro-cue onset. The upper panel shows contralateral and ipsilateral ERP waveforms; the lower panel shows the corresponding contralateral-minus-ipsilateral difference waveform. The ERP pattern showed a CDA-like posterior contralateral negativity after the retro-cue. **(c)** Posterior time-frequency lateralisation index relative to the second (non-cued) item and time-locked to the offset of the first memory probe. Alpha lateralisation (contralateral suppression) was already present shortly before completion of the first orientation report and was followed by a contralateral beta power increase. **(d)** Lateralised ERP activity time-locked to the offset of the first memory probe. No reliable lateralised ERP effect was observed in this second analysis window. Black contours in the time-frequency plots and black horizontal bars in the ERP difference plots indicate significant clusters. Dashed vertical lines indicate the relevant task events (first time window: memory array: -1050 ms, retro-cue: 0 ms; second time window: offset of first memory probe: 0 ms). Shaded areas in the ERP plots reflect 95% confidence intervals of the participant-level mean. Posterior channels PO7/8, P7/8, PO3/4 and P/6 were used for all analyses.

In the second time window, posterior alpha lateralisation associated with attentional re-orienting towards the item required for the second report was already present shortly before the end of the first orientation report. This stronger contralateral suppression of oscillatory power was followed by a contralateral increase in oscillatory power, with the strongest lateralised effect in the beta frequency range between approximately 20 and 30 Hz (Fig. 2c). No lateralised ERP effect was observed in this time window (Fig. 2d). Descriptively, the magnitudes of posterior alpha- and beta-power lateralisation, as well as the ERP lateralisation in the first time window, were highly similar to those observed in the subsequent blink-locked analyses (see Fig. 3), suggesting that both approaches captured these lateralised EEG effects to a comparable degree.

**Figure 3.**
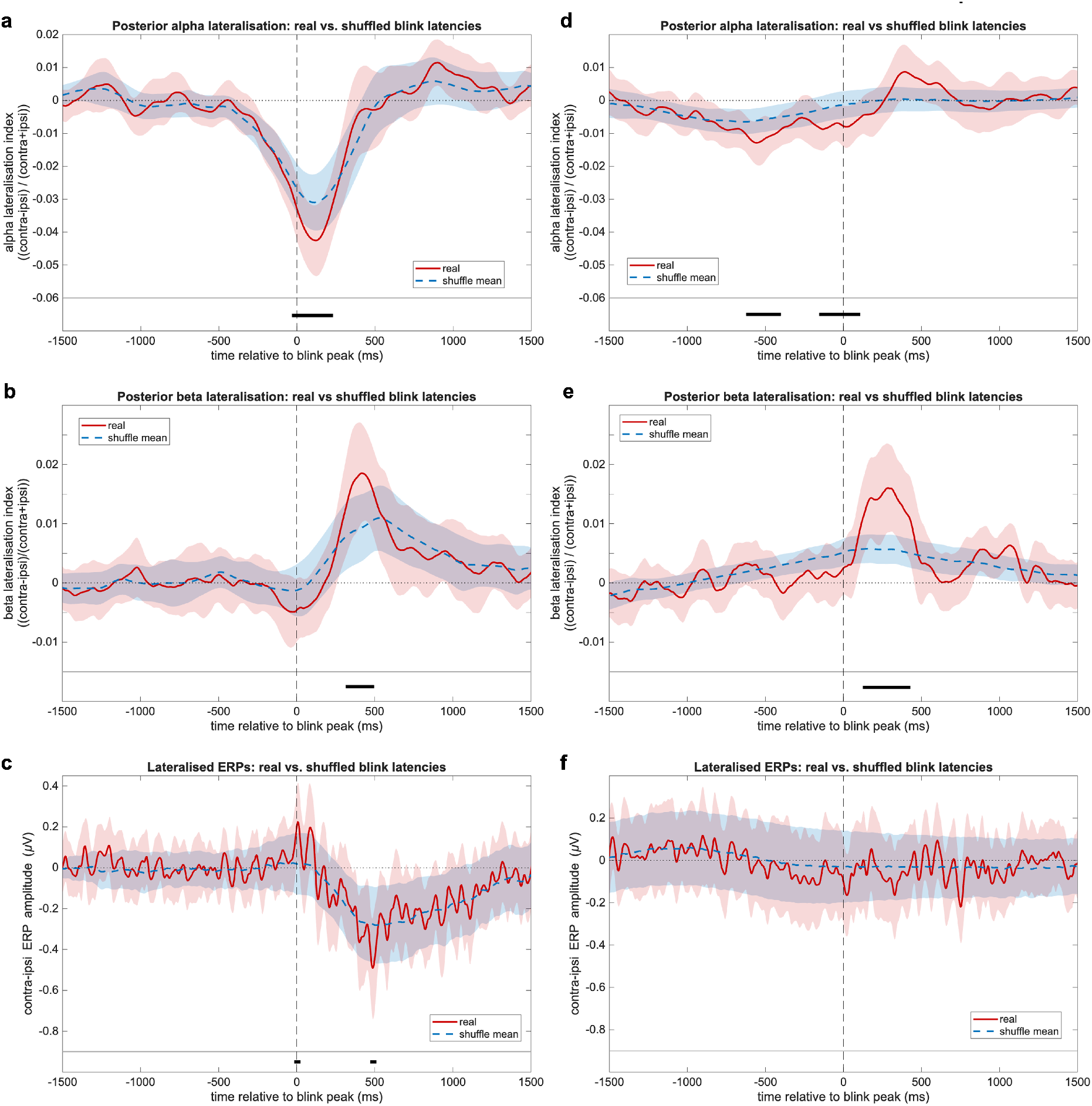
Blink-locked lateralised EEG activity in Experiment 1. Real blink-locked lateralisation waveforms are shown in red and latency-shuffled control waveforms (mean of all shuffles) in blue (dashed). Shaded areas indicate 95% confidence intervals. The left column shows time window 1, in which blinks were analysed after the retro-cue; the right column shows the second time window, in which blinks were analysed after completion of the first orientation report. Time zero denotes the blink peak. Black horizontal bars indicate significant time clusters from the real-vs-shuffled comparisons. **(a, d)** Posterior alpha lateralisation (8–13 Hz). In the retro-cue window, real blink timing was associated with stronger alpha lateralisation around blink peak, whereas in the second analysis window the real-vs-shuffled difference emerged also before the blink. **(b, e)** Posterior beta lateralisation (20–30 Hz). In both analysis windows, real blink timing was associated with a stronger contralateral beta increase shortly after the blink, showing a consistent post-blink profile across time windows. **(c, f)** Lateralised ERPs. Real blink timing revealed brief deviations from the shuffled estimate in the retro-cue window, including a transient contralateral positivity around blink peak and a brief contralateral negativity later in the trial, but no reliable ERP differences in the second analysis window.

For the blink-locked analyses, we examined EEG activity within the same two time windows, but time-locked to the peak of naturally occurring blinks, as measured from the ongoing EEG signal. Because blinks were not uniformly distributed across the trial (see Fig. 1c), we introduced a shuffling procedure based on 1000 permutations, which preserved the overall temporal distribution of blinks within the task while disrupting the specific alignment between individual blinks and EEG activity (see Methods for more details; see also supplementary Figure S4 for an illustration of how this shuffling procedure works with non-lateralised event-related potentials / ERPs). Comparing real blink-locked activity with this shuffled estimate thus allowed us to test whether blinks were temporally aligned with lateralised EEG activity beyond what would be expected from their overall timing within the task (see Fig. 3). Posterior alpha- and beta-power lateralisation were quantified by averaging oscillatory power across the 8–13 Hz and 20–30 Hz frequency ranges, respectively.

For posterior alpha power lateralisation relative to the location of the relevant working memory item, real blink timing revealed a reliable deviation from the shuffled estimate in both time windows. For blinks in the retro-cue interval, alpha lateralisation was stronger for real than shuffled blink times around the blink peak, indicating that blinks occurred in close temporal alignment with posterior alpha lateralisation (see Fig. 3a). In the second analysis window, in which blinks were referenced to the end of the first orientation report, the real-vs-shuffled alpha difference occurred already before the blink peak (see Fig. 3d). Thus, in both phases of the trial, true blink timing was associated with the timing of posterior alpha lateralisation, but the relative timing differed across phases. Results also revealed a lateralisation in posterior beta power in close succession to the alpha lateralisation, with higher contralateral than ipsilateral oscillatory power. Both after the retro-cue and after the first orientation report (see Fig. 3b and 3e), this posterior beta lateralisation was stronger for real than shuffled blink times shortly after the blink peak.

The lateralised ERP effects were comparatively brief and less consistent across the two time windows. In the retro-cue time window, real blink-locked activity showed short-lived deviations from the shuffled estimate, including a contralateral positivity around the blink peak and a brief difference in contralateral negativity from the shuffled estimate approximately 500 ms later. The latter effect occurred around the time at which the shuffled ERP waveform showed the strongest CDA-like contralateral negativity (see Fig. 3c). In the second analysis window, no reliable lateralised ERP differences between real and shuffled blink times were observed (see Fig. 3f).

Taken together, blink timing was most consistently aligned with posterior alpha- and beta-power lateralisation, with both measures differing reliably from the latency-shuffled estimates in both analysis windows.

### Experiment 2: Blink-frequency profiles track early and late selection in working memory

Experiment 2 tested whether the temporal profile of spontaneous blinking can reveal when selection among internal working memory representations is implemented. The task closely matched the temporal structure of Experiment 1, but introduced two retro-cue conditions that differed in when the relevant memory item could be selected for report. In the early-selection condition, a selective retro-cue indicated which of the two remembered items would have to be reported. Because retro-cues were 100% valid, participants could select the relevant internal representation immediately after the cue and maintain it in a prioritized state until report. In the late-selection condition, the retro-cue was neutral and did not indicate which item would become relevant. Selection therefore had to be postponed until the memory probe appeared, which indicated by its colour which item’s orientation had to be reported.

We first examined whether this manipulation affected behavioural performance and whether performance depended on the occurrence of a spontaneous blink between retro-cue and memory probe presentation (see Fig. 4). Orientation reports were more accurate in the early-selection than in the late-selection condition, confirming a robust benefit of being able to select the relevant memory item before the probe, F(1,27) = 249.494, *p* < 0.001, ηp² = 0.902, CI95 [-17.454°, -13.440°]. Across conditions, trials with a blink in the cue-to-probe interval did not differ reliably from trials without a blink, F(1,27) = 0.438, *p* = 0.514, ηp² = 0.016, CI95 [-0.939°, 0.481°]. Critically, however, the cue effect was qualified by an interaction between cue condition and blink occurrence, F(1,27) = 8.645, *p* = 0.007, ηp² = 0.243, CI95 [-2.814°, -0.501°]. Follow-up comparisons showed that blinks were associated with more accurate orientation reports in the early-selection condition, t(27) = -3.999, *p* < 0.001, dz = -0.756, CI95 [-1.6°, -0.515°], whereas no reliable blink-related performance difference was observed in the late-selection condition, t(27) = 1.047, *p* = 0.304, dz = 0.198, CI95 [-0.576°, 1.775°]. Thus, spontaneous blinks during working memory storage were not generally associated with better or worse working memory performance, but their behavioural relevance depended on whether they occurred during a period in which the relevant internal representation could already be selected prior to orientation report.

**Figure 4.**
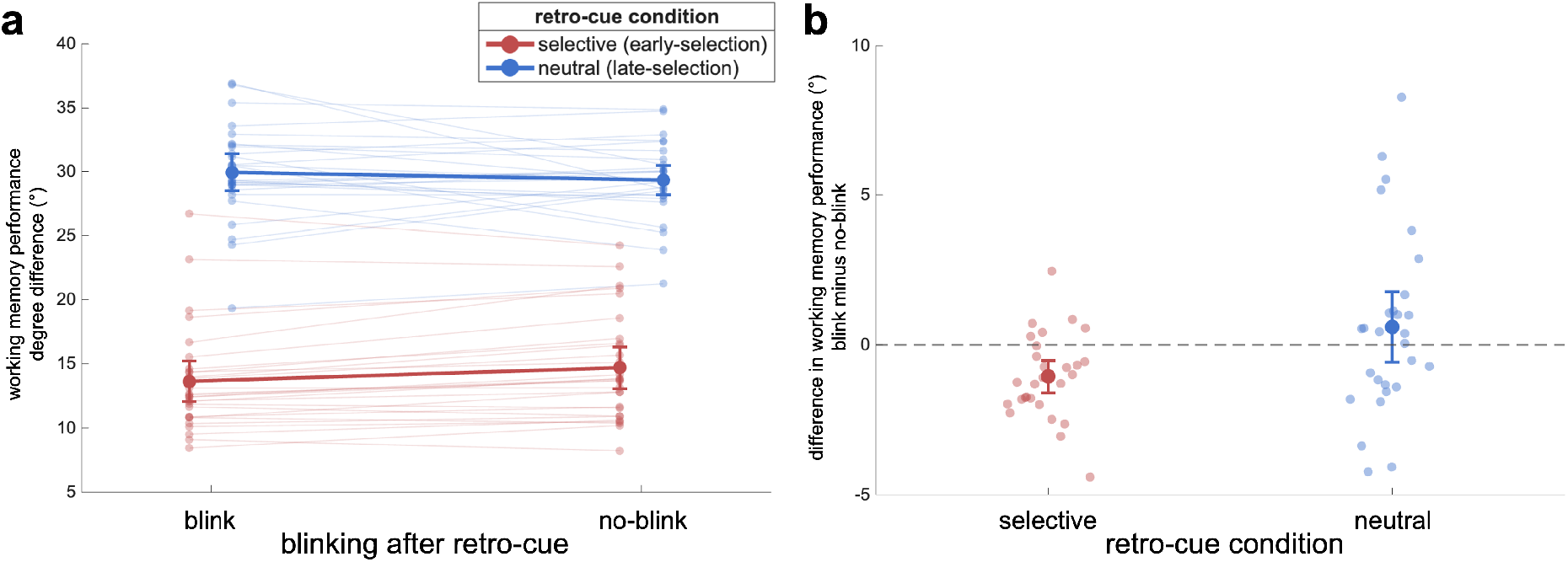
Effects of retro-cue condition and blink status on performance in Experiment 2. **(a)** Mean angular error / degree difference in orientation report for blink and no-blink trials, shown separately for selective (early-selection condition) and neutral cues (late-selection condition). Individual lines represent individual participants, and larger markers indicate group means. **(b)** Within-subject blink effects, computed as the difference between blink and no-blink trials for each participant, shown separately for selective and neutral cues. Negative values indicate smaller angular errors in blink trials than in no-blink trials, reflecting better performance when a blink occurred. Error bars indicate 95% confidence intervals around the respective group means.

We next asked whether the temporal profile of blink occurrence itself revealed when selection was implemented. For each participant, we calculated blink-frequency profiles in 20-ms bins relative to memory array onset. Because the first blink following a task event is most likely to carry chronometric information, we included only the first blink within each of three trial phases: memory array to retro-cue, retro-cue to memory probe, and memory probe to the end of the analysed epoch. Each trial could therefore contribute at most one blink per phase and three blinks overall. Condition differences in the resulting participant-level blink-frequency profiles were tested using a cluster-based permutation approach (see Fig. 5).

**Figure 5.**
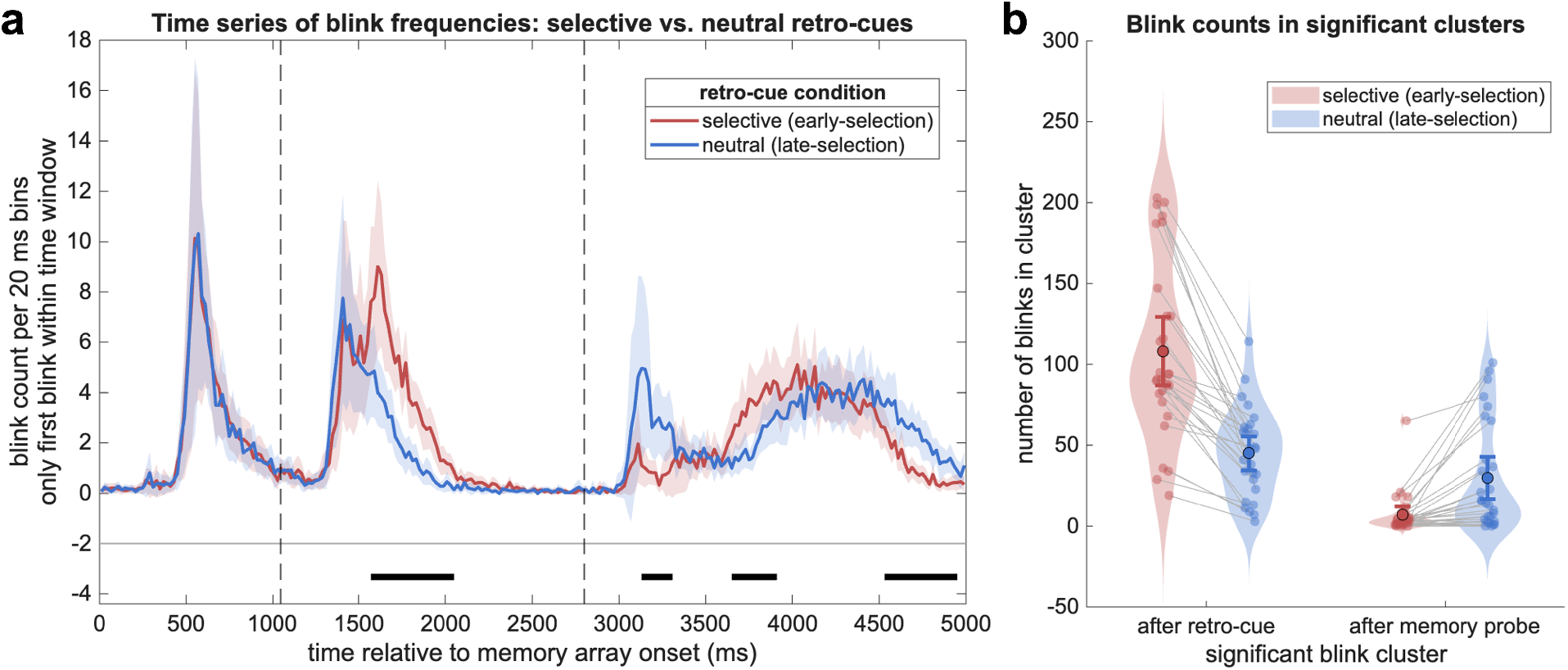
Blink frequency profiles across the trial in Experiment 2. **(a)** Time series of blink-frequency profiles for the selective and neutral retro-cue conditions, estimated in 20-ms bins relative to memory array onset. The profiles show a shared early post-cue increase, followed by condition-specific differences in later time windows (early- vs. late-selection increase). Vertical dashed lines reflect the onset of the retro-cue (1050 ms) and memory probe (2800 ms). Shaded areas reflect 95% confidence intervals of the participant-level mean**. (b)** Mean blink frequencies summarised across the significant selective-versus-neutral clusters identified by cluster-based permutation testing, separately for the post-cue and the first post-probe cluster. The post-cue cluster shows increased blink frequency in the selective condition, whereas the earliest post-probe cluster shows increased blink frequency in the neutral condition. Violin plots show the kernel density distributions of participant-level blink counts for the selective and neutral conditions. Individual participants are displayed as jittered points, with lines connecting paired observations across conditions, indicating that both clusters feature these differences consistently across participants. Larger circles indicate the group means, and error bars represent 95% confidence intervals around the respective group means.

The blink-frequency profiles showed both a condition-general and a condition-specific temporal structure. Shortly after the retro-cue, blink frequency increased in both the early- and late-selection conditions. Critically, the conditions diverged later in the cue-to-probe interval. In the early-selection condition, blink frequency showed a second post-cue increase that was absent, or markedly reduced, in the late-selection condition (t(sum) = 100.959, cluster *p* < 0.001). A complementary pattern emerged after memory-probe onset. In the late-selection condition, blink frequency increased shortly after the probe indicated which coloured item had to be reported, whereas this probe-locked increase was reduced in the early-selection condition (t(sum) = 30.273, cluster *p* = 0.002).

The key observation is therefore not simply that blink frequency increased after cue or after probe presentation, but that the timing of this increase shifted with the time at which selection of the relevant memory representation was possible. When the selective cue specified which orientation had to be selected before probe presentation, blink frequency increased during the later cue-to-probe interval. When the same cognitive operation had to be postponed until the probe identified the relevant orientation, a corresponding increase emerged only after probe onset.

The latency shift in blink frequency at the end of the trial (first cluster: t(sum) = 40.384, cluster *p* = 0.001; second cluster: t(sum) = 74.843, cluster *p* < 0.001) likely reflects condition differences in the completion of the manual orientation adjustment response (see Fig. 5a), since blinks are suppressed during visually guided action and occur more frequently once the response has been completed.^3^

Together, experiments 1 and 2 establish spontaneous blink timing as a behavioural chronometric tool for studying working memory. Experiment 1 linked blink timing in a working memory task to oscillatory correlates of attentional focusing (see Fig. 3). Experiment 2 showed that blinks were associated with higher performance when they occurred during the interval in which internal selection was possible and behaviourally relevant (see Fig. 4), and that blink-frequency profiles shifted in accordance with the experimentally manipulated timing of this interval (see Fig. 5).

## Discussion

### Blink timing is linked to attentional focusing in working memory

The present findings show that spontaneous blinks are systematically related to the temporal organisation of working memory processing. Experiment 1 helped constrain the interpretation of this relationship by linking spontaneous blink timing to task-defined lateralised EEG markers. Because these EEG measures were defined by the experimental manipulation, their lateralisation reflected task-related processing rather than activity related to the blink itself. Analysing these lateralised signals relative to blink timing therefore allowed us to ask whether eye blinks appeared temporally aligned with task-defined working memory operations.

Yet, how can we be sure that the observed relationship was driven by the trial-specific correspondence between blink timing and lateralised activity, rather than by the overall tendency of blinks to occur at particular moments within each phase of the trial? To address this question, we used a shuffling analysis in which blink latencies were reassigned across trials. The latency-shuffled controls preserved the average distribution of blink times relative to the relevant task event, that is, relative to the retro-cue in the first time window and relative to the end of the first orientation report in the second time window. Thus, the shuffled estimate retained the fact that blinks tended to occur at particular moments within each phase of the trial (see Fig. 1c). What was disrupted was the trial-specific mapping between the actual blink time and the neural signal from that same trial.

The clearest blink-specific temporal association was observed for posterior alpha power lateralisation. In retro-cue paradigms, this oscillatory effect can be interpreted as reflecting the orienting of attention towards an internal representation via its spatial contextual features, providing an indirect neural marker of internal attentional selection ^14,15,17–20^ (^21^ for a different perspective). Real blink timing revealed a clear deviation from the shuffled estimate in both analysis windows. In the retro-cue window, the alpha effect closely coincided with the blink peak (see Fig. 3a), whereas in the second delay interval the alpha effect emerged already before the blink peak (see Fig. 3d). This temporal pattern argues against a simple confound in which eye closure or blink execution produces stronger lateralised oscillatory activity: if the blink itself were sufficient to generate the stronger alpha power lateralisation, the effect should not already be present before the blink occurred. Instead, the results suggest that spontaneous blinks occurred at specific positions within the temporal sequence of task-defined attentional focusing. This relation should be understood statistically rather than deterministically: attentional focusing does not impose a fixed blink latency on individual trials. Rather, across trials, blink timing was systematically related to the unfolding of alpha power lateralisation, without requiring the stronger claim that the blink itself causes or modulates this oscillatory effect.

A related pattern was observed for posterior beta lateralisation, in the form of a contralateral beta increase shortly after the blink. Although this lateralised effect has received little to no attention in research on attentional focusing in working memory, comparable modulations are visible in previous work and may relate to post-selective adjustments following the spatial focusing of attention.^18,22^ Importantly, whereas the blink-locked alpha patterns revealed a stronger and more temporally consistent retro-cue-related alpha lateralisation, as expressed in a more pronounced and temporally concentrated alpha lateralisation around the blink, posterior beta lateralisation showed a more similar profile across both time windows. This argues against a simple reversal of the preceding alpha lateralisation effect in the beta frequency range and suggests that the contralateral increase in posterior beta power may be more closely linked to the blink itself, or to a stereotyped post-selective state around the blink. However, the present study was not designed to isolate the functional role of posterior beta power lateralisation. Since this effect has received little systematic attention so far, its precise functional significance requires further investigation.

The lateralised ERP results provided a further marker of spatially specific processing around blinks. No reliable ERP differences between real and shuffled blink timing were observed in the second time window (see Fig. 3f). In the time window after the retro-cue, however, real blink-locked activity showed brief deviations from the shuffled estimate (see Fig. 3c), indicating that phasic lateralised ERP activity relative to the cued location occurred in temporal association with the blinks. These effects were short-lived and are therefore more naturally interpreted as transient activity related to selection or attentional orienting than as evidence for a blink-related onset of sustained, storage-related ERP activity, such as the CDA.

Together, the posterior alpha lateralisation, posterior beta lateralisation, and lateralised ERP findings suggest that spontaneous blinks in the current task are specifically temporally embedded around the spatial focusing of attention within working memory. These presented parameters all relate to the cued location at a time when no sensory information is presented at this location. The results thus converge on the idea that the distribution of blink timings across trials can be informative about when spatial attention is focused on internal content, while leaving open the possibility that blinks covary with a working memory state that is closely coupled to, rather than identical with, the attentional process itself (see below). This raises the question of whether blinks are also behaviourally relevant when they occur during periods in which such attentional focusing within working memory is required.

### Blinks are behaviourally relevant when attentional focusing is engaged

The behavioural findings from Experiment 2 address this question by showing that blinks were linked to memory performance in a task-dependent manner. Blinks between retro-cue and memory probe were associated with more accurate orientation reports in the early-selection condition, where the selective retro-cue identified the relevant item before the memory probe appeared. In contrast, blink occurrence in the same interval was not associated with improved performance in the late-selection condition, where the neutral retro-cue did not yet specify which representation would become relevant (see Fig. 4). This finding does not rule out that blink rate can serve as a broader marker of cognitive effort or task demands; indeed, previous work has linked blink rate and blink-related measures to cognitive state variables, attentional resource allocation, and cognitive load,^1,23,24^ also in the context of working memory.^12^ Rather, the present results go beyond this more global interpretation by showing that the behavioural relevance of blinks depended on their position within the temporal structure of the task. In the present case, blinks were associated with better performance when they occurred during a period in which the relevant working memory representation was selected, stabilised, or brought into a report-ready state. The same interval in the late-selection condition did not yet afford these processes and featured no relation between the presence of blinks and working memory performance.

### Blink-frequency profiles as a behavioural chronometric tool

The central methodological contribution of the present study is that spontaneous blinks can be analysed not only as isolated events or average rates across phases of a trial, but as temporally resolved behavioural time courses (see Fig. 5).^11^ Blinking is a largely automatic behaviour. Although participants can intentionally produce or suppress blinks under explicit instruction, spontaneous blinks are typically distinguished from voluntary blinks and are not generated as deliberate task responses.^25,26^ Participants are therefore unlikely to have explicit access to the precise temporal pattern of their blinking across hundreds of trials, making blink-frequency profiles distinct from overt reports or deliberate response strategies. Their chronometric value emerges when these profiles are compared across experimental conditions that manipulate when a theoretically defined cognitive operation can occur. Condition-specific shifts in the temporal distribution of blinks can then indicate when the operation is likely to have occurred or when its consequences have established a new functional state for subsequent task processing (see Fig. 5a).

In working memory research, retro-cue designs provide a particularly suitable framework for this approach. In Experiment 2, participants encoded and stored the same type of memory representations in both conditions, but the point at which a single representation came to determine the subsequent report differed. In the early-selection condition, the retro-cue required to select the relevant representation during maintenance, creating an internal transition point at which the information guiding later behaviour could already be determined well before report.^27,28^ In the late-selection condition, this process was postponed until presentation of the memory probe. Both conditions showed an early post-cue increase, consistent with a processing consequence of the cue that was common to both conditions. However, the later condition-specific increase occurred after the retro-cue when it required attention to be focused on the relevant item, and after the probe when the target orientation could only then be identified (see Fig. 5). In the following section, we consider how this pattern can be interpreted in the context of the present task and in light of the converging evidence provided by the EEG-based findings from Experiment 1.

### A measurement logic for interpreting blink-frequency profiles

Blink-frequency profiles require the kind of interpretive caution familiar from other time-resolved measures such as event-related potentials (ERPs), whose functional interpretation depends on the experimental manipulation and the theoretical component logic.^29^ A peak in blink frequency does not necessarily mean that a particular cognitive operation is occurring at that moment, nor does the profile by itself identify the process that produced it. Relating such a time-resolved behavioural signal to the timing of latent cognitive operations therefore requires a suitable experimental design, a clear measurement framework, and, where possible, converging evidence from different approaches.

The present study illustrates this logic. Experiment 2 manipulated when the relevant representation had to be selected while otherwise keeping the temporal structure of the task closely matched. Selective and neutral trials occurred equally often, and all cues and probes were matched in luminance. Because selective cues and probes could be blue or orange, each individual colour occurred less often than the neutral grey stimulus. However, a visual-regularity account would predict comparable blink-frequency time courses following cue and probe, as the same regularity structure applied at both events and such effects are thought to arise rather automatically during early stimulus processing (for further details, see the supplementary results).^30^ Instead, the time courses differed markedly: the increase in blink frequency in the neutral condition peaked approximately 330 ms after memory probe presentation. The second increase in the selective condition peaked approximately 560 ms after retro-cue onset, overlapping with the period in which EEG and MEG markers of retrospective attentional focusing on relevant working memory representations typically emerge (see also Fig. 2).^17,18,31–33^ This observation also converged with the EEG results from Experiment 1, where blink timing was systematically aligned with posterior alpha lateralisation, a neural correlate of attentional focusing in working memory (see Fig. 3).

We thus conclude that, in the present tasks, blink-frequency modulations were chronometrically informative about internal attentional selection because this process coincided with a functional processing boundary. In the selective condition, identifying the relevant representation largely completed the processing required before report and allowed working memory to enter a temporarily stable, report-ready state, since the subsequent memory probe provided no further task-relevant information. In the neutral condition, this transition could not occur until it was specified which orientation had to be reported, and the blink-frequency increase shifted to after the memory probe. Accordingly, the observed pattern may reflect two complementary influences on blink timing: continued suppression while task-relevant information is still expected, as in the post-cue interval following the neutral retro-cue, and an increase in blink probability once the processing required to establish the relevant working memory representation has been completed. Thus, the frequency profiles need not directly index attentional selection itself. Instead, they may provide an indirect indication of how far working memory processing has progressed towards the requirements of a later task (here: the report of one specific orientation).

This interpretation is further supported by the association between blink occurrence and behavioural performance (see Fig. 4). Blinks were linked to more accurate reports when they occurred after the selective retro-cue had specified the relevant representation, but not during the corresponding interval in the neutral condition. This condition-specific association may indicate that blinks occurred more often on trials in which processing had already progressed sufficiently to establish the relevant representation for later report, whereas trials without a blink may more often have featured less advanced or incomplete target selection.

More broadly, the chronometric value of blink-frequency distributions may extend beyond working memory. Such distributions may provide temporal information whenever a cognitive operation marks a meaningful step towards completing a task. They should therefore not be viewed as a process-specific chronometric measure, but as a potential indicator of when latent processing reaches a functionally relevant stage before, or without, an overt behavioural response.

### Conclusion and outlook

Together, the present findings establish spontaneous blinks as a chronometric signal of latent cognitive processes on the level of working memory. Rather than merely confirming that blink rate varies with attentional state or processing demands, they show that the temporal distribution of blinks tracks a functional transition associated with internal attentional selection and the establishment of a report-ready state in working memory. Because blink timing is routinely recorded in EEG and eye-tracking studies, the same analytical approach could also be applied retrospectively to a wide range of existing datasets.

An important advantage of this approach is that blinks are easy to measure. In addition to laboratory EEG or eye-tracking studies, blink-frequency profiles could in principle be derived from camera-based recordings in online experiments.^34,35^ This would allow researchers to quantify an automatic behaviour that occurs naturally during task performance and can be measured without introducing an additional response requirement or interfering with task execution. Importantly, the detection of between-condition differences in aggregated blink-frequency profiles should not necessarily require millisecond-level precision, because condition effects can be estimated across trials and time bins, as illustrated by the 20-ms binning approach used here and by related considerations in mental chronometry.^36^ When combined with validated blink detection and theoretically constrained experimental designs, blink-frequency profiles therefore can provide a scalable and unobtrusive tool for studying the temporal organisation of cognition.

## Methods

### Participants

Thirty-two participants took part in Experiment 1 (mean age = 23.81 years, range = 19-29 years. Twenty-two participants identified as female and ten as male. Handedness (28 right-handers, 4 left-handers) was assessed by means of the Edinburgh Handedness Inventory (EHI).

Six participants were excluded from the EEG analyses because they featured fewer than 400 blinks in at least one of the two critical analysis windows across the 1200 experimental trials. This threshold was used to ensure sufficiently stable participant-level estimates for the blink-locked EEG analyses (see below). The sample used for EEG analyses thus comprised 26 participants (mean age = 23.65 years, range = 19-29 years, 19 females, 7 males, 23 right-handers, 3 left-handers). It is important to note that the analyses in Experiment 1 were re-analyses of data originally collected for a separate, preregistered study addressing the role of motor-planning processes in working memory. Accordingly, the sample size was determined for that original study rather than for the blink-locked EEG analyses reported here.

Thirty participants took part in Experiment 2 (mean age = 24.4 years, range = 19-30 years). Nineteen identified as female and eleven as male. Handedness (27 right-handers, 3 left-handers) was again assessed by means of the EHI. Two participants were excluded due to a very low frequency of eye blinks during the experiment (< 20 blinks following either the selective or neutral retro-cue), resulting in 28 datasets used for all analyses (mean age = 24.54 years, range = 20-30 years, 17 females, 11 males, 25 right-handers, 3 left-handers). As no comparable study had examined time-resolved blink-frequency profiles in relation to attentional selection in working memory, the sample size could not be based on a previously reported effect size. We therefore selected a target sample size comparable to those in closely related studies on attentional orienting in working memory. Studies using EEG, eye tracking, continuous report measures, and trial-rich retro-cue designs in this area commonly test samples in the range of approximately 23–31 participants.^20,37–41^ A target sample size of 30 participants therefore provided a sample size consistent with the most closely related literature.

All participants reported normal or corrected-to-normal vision, no history of neurological or psychiatric disorders, and intact colour vision as assessed using the Ishihara Test for Colour Blindness. Written informed consent was obtained prior to participation. Participants received €13 per hour or course credits in case of psychology students. The study was approved by the local ethics committee on 11 November 2024 and conducted in accordance with the ethical principles of the Declaration of Helsinki applicable at the time of ethical approval (approval number: 256). All participants provided written informed consent before participation.

### Apparatus, stimuli, and procedure

The experiments were carried out in a dimly lit, electrically shielded room using a ViSaGe MKII Stimulus Generator (Cambridge Research Systems, Rochester, UK). Visual stimuli were displayed on a 22-inch CRT monitor with a resolution of 1024 × 768 pixels and a refresh rate of 100 Hz. For Experiment 1, viewing distance was ∼100 cm. Due to the use of an eye-tracker, participants completed Experiment 2 with their head stabilised in a chin rest, and viewing distance was set to 50 cm. The eye-tracking data were used primarily to validate the reliability of blink latencies derived from the EEG signal, as described in the supplementary results section. No further eye-tracking analyses are reported here. The experimental paradigm was implemented in Lazarus IDE using Free Pascal.

Prior to the experiments, participants completed a demographic questionnaire assessing age, sex, visual correction, medication use, and any history of neurological or psychiatric disorders, to verify inclusion criteria (age: 18-30 years, no history of psychiatric or neurological disorders). They also filled out the EHI and were tested for colour blindness by means of the Ishihara Test. Before the main experiments, participants completed 80 training trials for Experiment 1 and 40 trials in Experiment 2. These training sessions represented a shortened version of the full experimental procedure and included all experimental conditions. The training served to familiarize participants with the task requirements and response procedures.

### Experiment 1

Each experimental trial began with the presentation of a central fixation point (diameter: 0.33° visual angle), which remained on the screen throughout the trial. With a jittered delay between 800 and 1200 ms (randomised), a memory array consisting of two oriented stimuli was presented for 250 ms. The stimuli were two bars (3.5° x 0.35° visual angle) with one bar presented in orange (CIE1931 color space: x = 0.500, y = 0.432, Y = 40 cd/m²) and the other presented in blue (x = 0.193, y = 0.221, Y = 40 cd/m²) on a dark-grey background (x = 0.291, y = 0.310, Y = 15). Stimulus colours were assigned randomly to the side of presentation, with the constraint that colour-side combinations occurred equally often across trials. The orientations were sampled from a continuous range between 1° and 180°, with near-vertical orientations excluded (+/- 5° from vertical). To ensure a balanced representation of orientation relationships, trials were constructed such that there was an equal number of pairs in four categories: (i) both bars tilted leftward, (ii) both bars tilted rightward, (iii) left bar leftward and right bar rightward, and (iv) left bar rightward and right bar leftward. This balancing was included to allow the contribution of prospective motor representations to working memory performance to be assessed, although this was not the central research question of the present study.

After a brief retention interval of 800 ms, a retro-cue was presented for 250 ms and indicated by colour (blue vs. orange) which of the two items was relevant for the first report. The retro-cue was followed by another delay period of 1500 ms. Upon presentation of the first memory probe (a grey bar: x = 0.291, y = 0.310, Y = 40), participants reported the orientation of the cued item by rotating the probe stimulus using either a left (counterclockwise rotation) or right (clockwise rotation) button press on a standard keyboard until it reached the desired orientation. Participants were given 2000 ms to complete this task (see Fig. 1a).

Although this aspect was not central to the present analytical approach, which focused on lateralised EEG effects relative to the presentation side of the relevant information held in working memory, there were two different kinds of memory probes: There were informative memory probes, meaning that the starting orientation of the memory probe was always vertical (0°) and participants had to use either the left (counterclockwise rotation) or right (clockwise rotation) button to reach the cued orientation, because memory probe stimuli could only be rotated for up to 90°. Furthermore, there were random memory probes with random starting orientation that could be rotated either clockwise or counterclockwise to reach the cued orientation. These probe types were assigned to either the cued item report or non-cued item report (i.e., reported second) in a block-wise fashion, so that both probe types appeared in each trial. These manipulations were part of a separate research project on prospective motor representations in working memory.

After completing the first report (defined by button release, the maximum rotation reached in the informative probe condition, or the offset of the memory probe stimulus after 2000 ms), a second delay period of 1500 ms followed, during which participants could re-focus attention on the previously non-cued item for the subsequent orientation report. The second orientation report also had to be completed within 2000 ms and proceeded to ITI (800-1200 ms) either upon button release, upon reaching maximum rotation (informative probe condition), or after those 2000 ms had elapsed.

The experiment consisted of 1200 trials, divided into 12 blocks of 100 trials each. The probe conditions (informative vs. random) were assigned to either the first or second orientation report in a block-wise fashion, with block order counterbalanced across participants. Fixation was to be maintained throughout each trial. There was no instruction at all concerning eye-blinks, which ensures that blinking behaviour was not controlled based on a secondary task (like being allowed to only blink during specific phases of a trial).

### Experiment 2

Experiment 2 matched Experiment 1 in timing up to the report of the cued orientation.

Stimulus size, eccentricity and spacing in degrees of visual angle, as well as luminance and colour values, were also kept identical across experiments. However, the exact procedure differed between experiments. Whereas both orientations had to be reported sequentially in Experiment 1, only the cued orientation had to be reported in Experiment 2 (by adjusting the orientation of a memory probe to match the cued item). The relevant orientation was indicated either by a selective retro-cue or, in the late-selection condition, by the colour of the memory probe. Accordingly, Experiment 2 ended after the single orientation report and did not include a second memory probe for the previously irrelevant orientation. Thus, attentional selection could occur either during the retro-cue interval (early-selection) or only at the time of the memory probe (late-selection). In the late-selection condition, the retro-cue was replaced by an equally sized and equally bright grey circle (x = 0.291, y = 0.310, Y = 40), so that it provided no information about which item would later become relevant. Memory probe starting orientation was always vertical (corresponding to the informative probe condition of Experiment 1). The experiment consisted of 800 trials, divided into 10 blocks of 80 trials each. Half of the trials included a selective retro-cue and half a neutral retro-cue. Within each condition, the left and right orientation was relevant for report in 200 trials each. The relevant orientation was indicated by blue or orange with equal probability, either by the retro-cue in the selective condition or by the later memory probe in the neutral condition. Again, there was no instruction at all concerning eye-blinks.

### Behavioural analyses

#### Experiment 1

Task accuracy was measured by the circular absolute degree difference between the to-be-reported and the set orientation for each report position (first vs. second report). For the informative memory probe conditions, these parameters were calculated only across those trials in which participants used the correct hand for the rotation response. In the random probe condition, probe starting orientations varied across trials, allowing multiple possible rotation paths. Thus, to make the way of responding comparable to the informative condition, only trials in which participants chose the shortest rotation direction were included when calculating angular report error. Working memory performance was then compared between the two report positions (first vs. second orientation report) by means of a two-sided within-subject t-test.

Eye-blinks were considered in two time-windows: The first time-window started after retro-cue presentation and lasted until presentation of the first memory probe. The second time-window started after offset of the first memory probe (when the first orientation had been reported or max. available time or rotation was reached) and lasted until presentation of the second memory probe (see Fig. 1c). Details about eye-blink detection from the continuous EEG signal are provided in the section concerning EEG data pre-processing.

#### Experiment 2

Task accuracy was measured by the circular absolute difference between the to-be-reported and the set orientation. To test whether blink occurrence during the post-cue interval was related to working memory performance, trials were classified according to whether a blink occurred after retro-cue presentation and before memory probe onset. Mean report error was then analysed using a 2 × 2 repeated-measures ANOVA with the factors blink occurrence (blink vs. no blink) and cue condition (selective/early-selection vs. neutral/late-selection). Details about eye-blink detection from the continuous EEG signal were identical between Experiments 1 and 2 and are provided in the section concerning EEG data pre-processing (see below).

In a second analysis step, we analysed blink-frequency profiles across the trial, from memory array presentation to 5000 ms thereafter, to test whether the temporal distribution of blinks differed between retro-cue conditions. For each participant and condition, blink frequency was calculated in 20-ms bins, reflecting the number of blinks within each bin across trials.

Importantly, only the first blink within each phase of the trial (after memory array, after retro-cue, after memory probe) was considered. These blink-frequency profiles were compared between retro-cue conditions using cluster-based permutation testing across the 20-ms time bins, to control for multiple comparisons across the time series. For each time bin, the within-subject difference between conditions was tested against zero using a two-sided t-test. Adjacent bins exceeding the cluster-forming threshold of *p* < .05 and showing the same effect direction were grouped into clusters separately. Within each cluster, the signed *t*-values were summed to quantify cluster strength. Statistical significance was assessed using a sign-flipping procedure with 10000 permutations. For each permutation, the condition-differences were randomly sign-flipped across participants, and the largest absolute cluster-level sum of *t*-values was retained to form the null distribution. An observed cluster was considered significant if the absolute value of its summed *t*-values exceeded the 95th percentile of this permutation distribution. For all experiments, effect sizes based on within-subjects t-tests are reported as Cohen’s *d_z_*. For ANOVAs, effect sizes are reported as partial eta squared (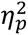). Cohen’s *d* for paired comparisons was calculated by dividing the mean of the participant-wise difference scores by their standard deviation. Partial eta squared was calculated from the corresponding *F*statistic and its degrees of freedom as 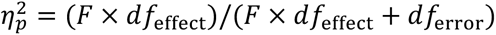

### Experiments 1 and 2: Pre-processing of EEG data

EEG was recorded from 64 passive Ag/AgCl electrodes mounted in an Easycap electrode cap and arranged according to the internation 10-10 system. Signals were acquired at a sampling rate of 1000 Hz using a NeurOne Tesla AC amplifier (Bittium Biosignals), with an online low-pass filter at 250 Hz. FCz served as the online reference and AFz as the ground electrode. All analyses were conducted in MATLAB (R2025b; MathWorks) and with the help of the EEGLAB toolbox (v2025.0.0).

EEG data were used differently across the two experiments. Experiment 1 included a substantially larger number of trials required for the shuffling approach based on EEG lateralisations. It was therefore used for the main analyses of task-related lateralised EEG activity. In Experiment 2, the EEG signal was used primarily to determine the timing of blink peaks. Simultaneously with the EEG recording in Experiment 2, eye position was recorded using an EyeLink 1000plus eye tracker (SR Research) at a sampling rate of 1000 Hz, tracking the horizontal and vertical position of both eyes. In the present investigation, these data were used to validate that blink latencies could be reliably derived from the EEG signal alone (see supplementary results; Fig. S1 and S2). EEG preprocessing was identical to Experiment 1.

Continuous data were band-pass filtered between 0.1 and 40 Hz using zero-phase finite impulse response (FIR) filters as implemented in EEGLAB (pop_eegfiltnew; default settings with Hamming window and automatically determined filter order). Noisy channels were identified using an automated kurtosis-based procedure (pop_rejchan) applied to all electrodes except a predefined set of frontal ocular channels (Fp1, Fp2, AF3, AF4, AF7, AF8) to avoid bias from eye-related activity. Specifically, channels were rejected if their normalized kurtosis exceeded 15 standard deviations from the mean across channels, indicating abnormally peaked or heavy-tailed signal distributions. Identified channels were removed and later reconstructed. The data were subsequently re-referenced to the common average. This dataset (filtered, channel-cleaned, and re-referenced at the original sampling rate) was retained as a reference dataset.

For ICA decomposition, a copy of the data was down sampled to 200 Hz and high-pass filtered at 1 Hz (same FIR filter design) to improve decomposition quality. Data were segmented into epochs ranging from −1 to 6 s relative to memory array onset and screened using automated trial rejection as implemented in EEGLAB (with default settings). Independent component analysis (ICA) was then performed using the extended Infomax algorithm with PCA-based rank reduction (number of components = number of channels − 1). ICs were classified using ICLabel.^42^ Components exceeding a probability threshold of 0.4 for the eye artifact class were marked for removal. ICA weights were then transferred back to the reference dataset, thereby preserving the original temporal resolution for subsequent analyses.

For both Experiments 1 and 2, eye blinks were subsequently detected on these continuous data prior to component rejection, using the BLINKER algorithm.^43^ Detected blink peaks were inserted as events into the EEG data. For the blink-locked EEG analyses in Experiment 1 (see below), only the first blinks occurring within a latency window of 150–1500 ms relative to the retro-cue (first time-window) and relative to the offset of the first memory probe (second time-window) were re-labelled as task-relevant blink events. These restricted time windows were used to exclude blinks overlapping with the presentation of critical task events. After blink detection, the identified eye-related independent components were removed from the continuous signal.

Then, previously rejected channels were reconstructed using spherical interpolation. Finally, continuous EEG data were epoched from -1000 to 5000 ms relative to memory-array onset for the analyses of the first time window, which encompassed the period before and after the retro-cue up to the first memory probe. For the analyses of the second time window, data were epoched from -1000 to 7800 ms relative to memory array onset, thereby including the later trial period after completion of the first orientation report. The extended epochs were essential for the later analyses of blink-locked lateralised EEG effects. These time-series data were baseline-corrected using the 200-ms interval preceding memory array onset. Artefact rejection was then performed separately for the two sets of epochs using EEGLAB’s amplitude-threshold rejection procedure. Epochs in which any EEG channel exceeded ±400 µV were rejected from further analyses.

### Experiment 1: EEG analyses

#### Time-frequency decomposition

Time-frequency oscillatory power was estimated on the epoched EEG data using complex Morlet wavelet convolution. Before decomposition, data were resampled to 250 Hz. Power was computed for 26 logarithmically spaced frequencies between 4 and 30 Hz. The number of wavelet cycles increased logarithmically across frequencies from 3 to 11.25 cycles, providing a compromise between temporal resolution at lower frequencies and frequency resolution at higher frequencies. For the first and second analysis time windows, decomposition was performed on the epochs from −1000 to 5000 ms and −1000 to 7800 ms relative to memory array onset, respectively.

For each frequency, a complex Morlet wavelet was generated and multiplied pointwise with the single-trial EEG signal in the frequency domain. Multiplication was performed separately for each channel and epoch, and power was obtained as the squared magnitude of the resulting complex convolution signal. The resulting time–frequency data were stored as raw power values for each channel, time point, frequency, and epoch. No baseline correction was applied.

The edge exclusion window was defined from the nominal temporal extent of the wavelet. For each frequency, this extent was calculated as the number of cycles divided by frequency, and half of this duration was excluded at each epoch boundary. To ensure a common valid output time range across frequencies, the exclusion window was defined by the largest half-window, corresponding to the lowest frequency (376 ms).

### Lateralised EEG effects

The main analytical approach focused on lateralised EEG effects. Under the assumption of a well-controlled experimental design, such effects provide a direct index of spatially selective processing of relevant working memory content. We analysed these effects in both stimulus-locked and blink-locked data. For the stimulus-locked oscillatory analyses (see Fig. 2), lateralisation was examined across the full frequency range from 4 to 30 Hz. This range typically captures stronger contralateral suppression of alpha power, a well-established marker of spatial attentional orienting, also in the context of retro-cues in a visuo-spatial working memory task.

We additionally observed a less commonly described contralateral increase of oscillatory power in the beta frequency range. Based on these frequency-specific patterns, the subsequent blink-locked analyses were restricted to average power in the alpha (∼8–13 Hz; exact frequencies: 8.26 Hz - 13.4 Hz) and beta (∼20–30 Hz; exact frequencies: 20.05 Hz – 30 Hz) ranges.

The lateralisations in oscillatory power were calculated from a posterior electrode region of interest, comprising PO7/PO8, P7/P8, PO3/PO4, and P5/P6.^14,19^ For each participant, they were computed on non-baselined power values (raw power, µV²) as a lateralisation index, defined as the difference between contralateral and ipsilateral power divided by their sum, relative to the side of the relevant working memory item. Contralateral and ipsilateral oscillatory power were calculated by averaging activity across the corresponding electrodes within the posterior region of interest. In the first time window, this lateralisation was defined relative to the item indicated by the colour of the retro-cue. In the second time window, it was defined relative to the item that remained relevant for the second report.

Second, we analysed lateralised ERPs. ERP lateralisation was calculated over the same posterior electrode region of interest after baseline correction, using the contralateral-minus-ipsilateral difference rather than the normalised lateralisation index applied to oscillatory power. Phasic posterior ERP lateralisation can reflect attentional selection processes, as described for components such as the N2pc or posterior contralateral negativity.^44,45^ In addition, more sustained posterior contralateral negativity can index the active maintenance of relevant visuo-spatial information in working memory, commonly referred to as the contralateral delay activity (CDA^16^). The ERP analyses therefore allowed us to test whether blinks were temporally aligned with transient selection-related ERP activity or with the onset of a more sustained lateralisation related to active storage of visuo-spatial information in working memory.

For the stimulus-locked analyses, we used the same sign-flipping cluster-based permutation procedure as for the blink-frequency time courses in Experiment 2 (see above), applied across adjacent time points for the ERP data and adjacent time x frequency points for oscillatory power.

### Shuffling approach for assessing blink-locked lateralised EEG effects

To assess whether the observed blink-locked EEG lateralisations were specifically related to the true timing of blinks, we used a latency-shuffled control analysis (see Fig. 3). Importantly, the single-trial time-frequency decomposition was computed only once per participant and was not recomputed for each shuffle. Instead, only the assignment of blink latencies to trials was shuffled within each analysis window. Shuffling was performed relative to the relevant task event: in the first analysis window, blink latencies were shuffled relative to the retro-cue, whereas in the second analysis window, blink latencies were shuffled relative to the end of the first orientation report. Additionally, shuffling was performed separately for trials with left- and right-sided relevant working memory items to preserve any systematic differences in average blink latency between these two conditions. This procedure preserved the overall distribution of blink times within each time window, while disrupting the trial-specific mapping between the actual blink time and the neural signal from the same trial.

For each participant and analysis window, blink-locked lateralisation was recomputed for each shuffled latency assignment using the same single-trial EEG or time-frequency data. To ensure sufficiently stable participant-level estimates across the large number of shuffled assignments, these analyses were restricted to datasets contributing more than 400 blinks meeting the inclusion criteria defined above in each of the two time windows. Six of the 32 datasets did not meet this criterion and were excluded. The resulting shuffled waveforms therefore provided participant-specific estimates of the lateralised signal expected when the temporal relationship between the true blink time and the corresponding neural activity was disrupted.

We then computed the difference between the real blink-locked lateralisation and the participant-specific mean of the shuffled estimates. At each time point, paired-samples t-statistics were computed across participants for this real-minus-shuffled difference. We first identified time points at which this difference was unlikely to be zero using a two-sided threshold of p < 0.05. Adjacent significant time points were then grouped into positive and negative clusters separately. Within each cluster, the signed t-values were summed to quantify its strength.

To obtain the permutation null distribution, we performed 10000 permutations based on the shuffled data. For each permutation and participant, one shuffled estimate was randomly selected and treated as a pseudo-real blink-locked estimate. This pseudo-real estimate was then compared with the mean of the remaining shuffled estimates from the same participant. The resulting differences were entered into the same group-level cluster procedure as the observed real-minus-shuffled difference. For each permutation, the largest absolute cluster-level sum of t-values across the blink-locked time series was retained. This yielded a null distribution of maximum cluster statistics expected if blink timing was not specifically aligned with lateralised EEG activity. An observed cluster in lateralisations based on the real timing of the blinks was considered significant if the absolute of its summed t-values exceeded the 95th percentile of this permutation-based distribution.

## Acknowledgements

We thank Tobias Blanke for the invaluable technical support and for programming the experiments. We also thank Rama Alnabulsy for her outstanding assistance with participant recruitment and data collection.

## Supplementary Results

### 1. Eye tracking and EEG-based blink detection in Experiment 2

The eye-tracking signals were recorded via the EyeLink 1000plus analogue outputs, digitized by the EEG acquisition system, and stored as additional channels in the EEG file. We did not convert them back into calibrated screen coordinates, since the present analysis used these signals only to assess the temporal alignment between EEG-derived blink triggers and the blink-related eye-position artifact (i.e., the signal changes surrounding the temporary loss of pupil tracking).

Both EEG and eye-tracking data were epoched around memory-array onset (-1000 to 3500 ms). Trials with abnormal pre-stimulus fixation were rejected using an outlier procedure. For each trial and eye-position channel, the median signal in the 500 ms interval preceding memory array onset was computed. These trial-level fixation estimates were then z-scored across trials separately for each channel, using the median and median absolute deviation. A trial was rejected if the absolute z-score exceeded 4 in any of the eye-position channels. The remaining eye-position data were baseline-corrected by subtracting the median pre-stimulus eye position separately for each channel and trial.

These data were then epoched around the EEG-derived blink trigger. Within each blink-locked epoch, the blink-related eye-position artifact was characterised separately for the horizontal and vertical eye-position channels. For each channel, the algorithm identified the strongest blink-related signal changes before and after the EEG-derived blink trigger. These changes reflect the transition into and out of the blink, that is, the artifact phases associated with eye closure and reopening. The interval between these transitions was used to estimate an inner blink phase. During this phase, loss of pupil tracking caused the recorded eye-position signal to approach zero. Because the data had been baseline-corrected, this signal loss appeared as a marked negative deflection. Thus, the negative excursion reflects temporary loss of the tracking signal during eye closure rather than an actual change in gaze position.

The broader blink-artifact interval was defined by extending the detected blink-related edges to nearby periods of stable velocity and adding a 20-ms temporal padding before the estimated onset and after the estimated offset.

**Figure S1.**
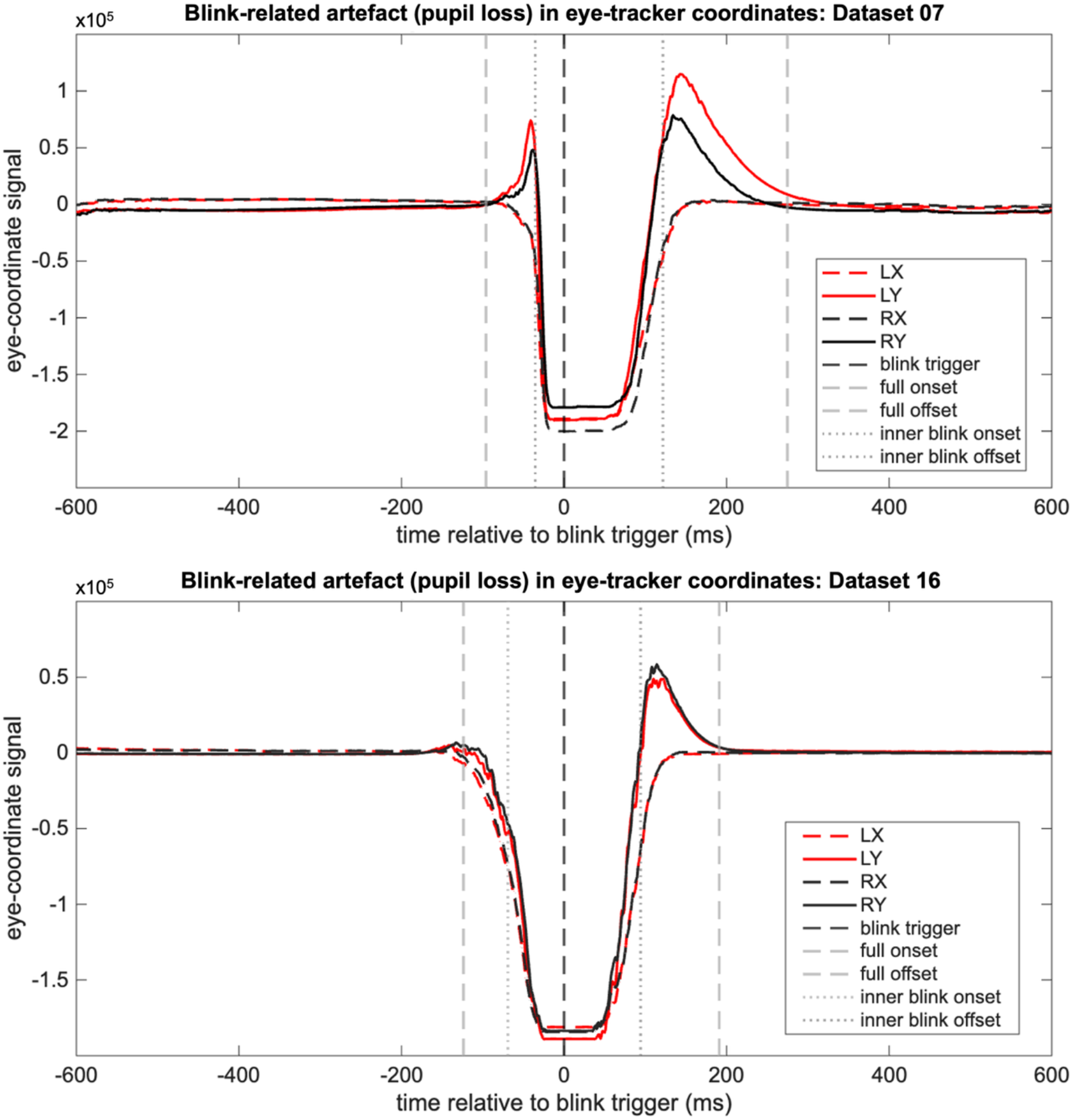
Exemplary eye-position signal for datasets 7 and 16. Time-point zero reflects the EEG marker set with the help of the BLINKER toolbox for EEGLAB. The negative values reflect the average phase of signal loss during eye-tracking for each dataset, not actual changes in gaze position. The grey dashed vertical lines reflect the average onset and offset of the broader blink artefact, while the dotted grey lines reflect the inner phase of the eye blink as an estimated equivalent to average eye-closure time.

These onset- and offset-times were averaged across epochs separately for each eye-position channel, yielding dataset-level descriptive estimates of the onset and offset of the full blink-related eye-position artifact and of the inner signal-loss phase, the latter providing an approximate measure of eye-closure duration. Figure S1 shows exemplary results from two datasets and indicates that the blink peak estimated from the EEG occurred closer to the beginning of the eye-closure interval.

**Figure S2.**
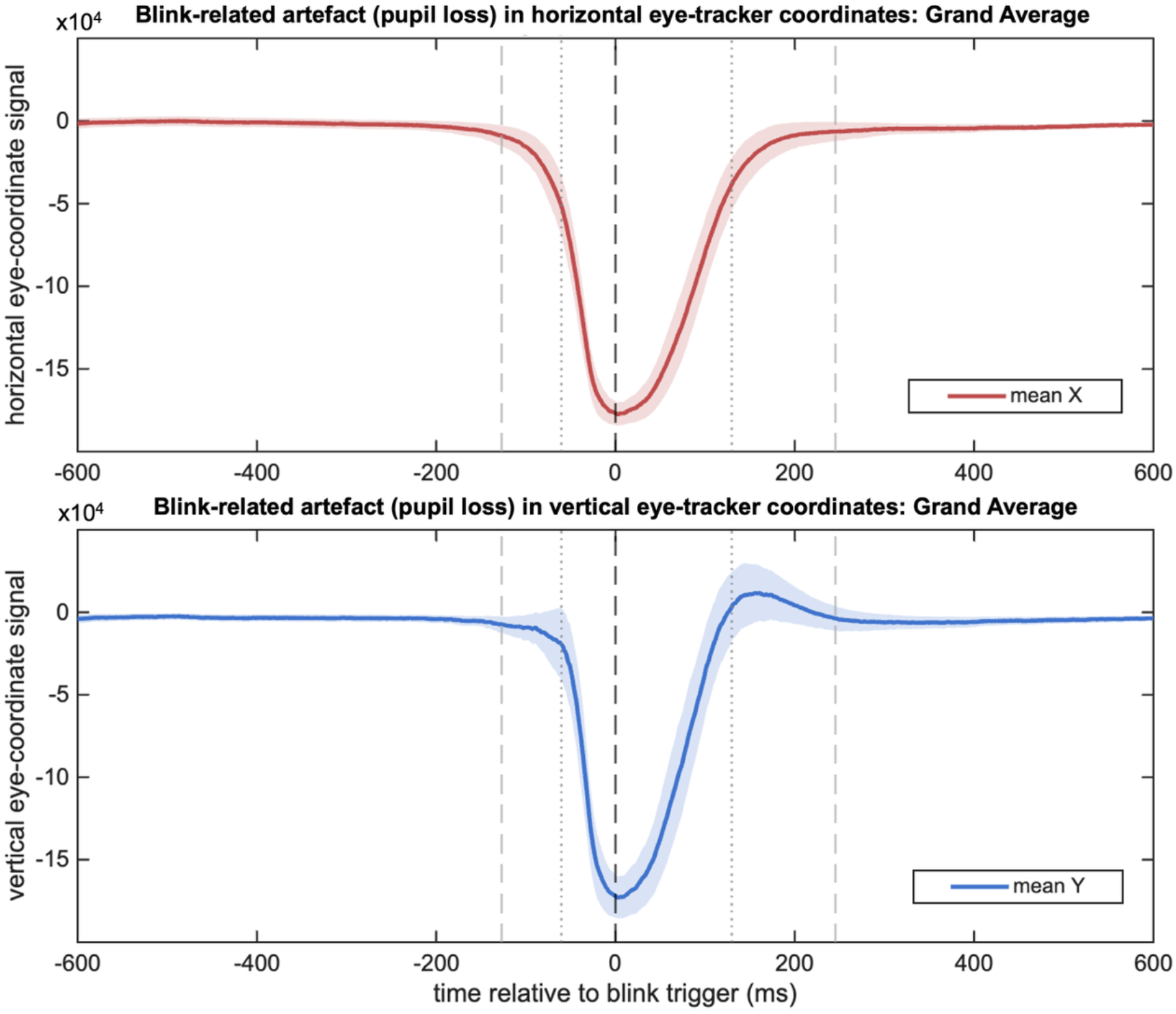
Average blink-locked eye-position traces across included datasets. Grand-average eye-position signals recorded from the eye-tracker channels embedded in the EEG recording, time-locked to blink triggers identified in the EEG signal. Horizontal and vertical eye-position traces were computed by averaging the corresponding left- and right-eye channels within each participant and then averaging across included participants. The negative values reflect the average phase of signal loss during eye-tracking across datasets, not actual changes in gaze position. Shaded areas indicate 95% confidence intervals around the mean across participants. The black dashed vertical line marks the EEG-derived blink trigger. Gey dashed vertical lines indicate the mean onset and offset of the full blink-related eye-position artifact, and grey dotted vertical lines indicate the mean onset and offset of the inner blink phase, estimating average eye-closure time.

For the group-level validation (see Fig. S2), blink-locked eye-position traces were averaged across blink epochs within each dataset and then across datasets. Only datasets included in the blink-related analyses within the manuscript entered this grand average (n=28). Horizontal and vertical eye-position signals were obtained by averaging the corresponding left- and right-eye coordinate channels within each participant. As in the single-dataset examples, temporary loss of pupil tracking during eye closure caused the recorded eye-position signal to approach zero. Because the data had been baseline-corrected, this signal loss appeared as a pronounced negative deflection rather than reflecting an actual change in gaze position. The signal-loss phase does not appear as a sharp, rectangular plateau in the grand average because its onset, duration, and offset vary across blinks and datasets.

Averaging across these temporally variable signal-loss intervals therefore smooths their boundaries and produces a gradual deflection rather than abrupt edges. At the group level, the EEG-derived blink trigger occurred close to the peak of this negative deflection, indicating that it closely marked the point at which eye closure was most consistently observed across blinks and datasets. This supports the use of the peak of the blink artifact in the EEG signal as the temporal reference for the blink-locked analyses reported in the manuscript.

### 2. The temporal relationship between cue-related and probe-related blink frequencies

To determine whether the condition difference in blink timing after the retro-cue could account for the subsequent blink response after the memory probe, we conducted an additional trial-level analysis separately for the selective- and neutral-cue conditions. The analysis was designed to test whether the comparatively later post-cue blinks in the selective condition were followed by a reduced post-probe blink response, as would be expected if the two effects were related through a simple blink-recency or refractory mechanism.

As with the analyses provided in the manuscript, 28 participants with at least 20 blinks in the post-cue period for both selective and neutral retro-cue were included in the analyses. Within each participant and cue condition, trials were classified according to the first blink occurring between retro-cue onset and memory-probe onset. Trials containing a post-cue blink were divided using a participant-specific median split. Trials in which the first post-cue blink occurred before the participant’s median latency were assigned to the early-blink category, whereas trials with a first post-cue blink at or after the median were assigned to the late-blink category. If the number of post-cue blink trials was odd, the central trial was excluded to obtain equally sized early- and late-blink groups. The classification therefore reflected the relative timing of post-cue blinking within each participant and condition rather than absolute latency differences between participants. Then blinks in the first (from memory array to retro-cue onset) and third phase of the trial(from memory probe onset onward), were categorized separately within each participant and retro-cue condition according to whether they appeared as first blink in the trials categorized into early vs. late conditions based on post-cue blink latency. Within these resulting trial subsets, blinks were counted in 20-ms bins. For visualization, the corresponding time courses were also plotted for the second, post-cue phase. Paired two-sided cluster-based permutation tests with sign-flipping across participants were applied separately within the selective and neutral retro-cue conditions to compare the time-resolved blink frequencies between trials classified as early versus late based on blink latency in the post-cue phase. Because the blink rates in these phases depend on the overall incidence of post-cue blinks in the second phase, they are condition-dependent and should be interpreted only for the early-versus-late comparison within the retro-cue conditions. For visualisation, participant-level distributions were smoothed using a 40 ms moving average window.

As also evident from the temporal profiles reported in the main manuscript, the selective retro-cue condition showed a double-peaked blink frequency profile in both the early- and late-classified trial subsets. Importantly, the later of these two peaks (i.e., the component that primarily distinguished the selective from the neutral retro-cue condition) occurred approximately 200 ms later as compared to the selective vs. neutral retro-cue effect relative to memory probe onset. This temporal discrepancy makes it highly unlikely that the cue-related and probe-related modulations reflect the same underlying event-locked response to visual feature regularity or predictability. If both effects arose from a common response triggered by the respective visual events, their temporal profiles relative to event onset should be closely aligned.

**Figure S3.**
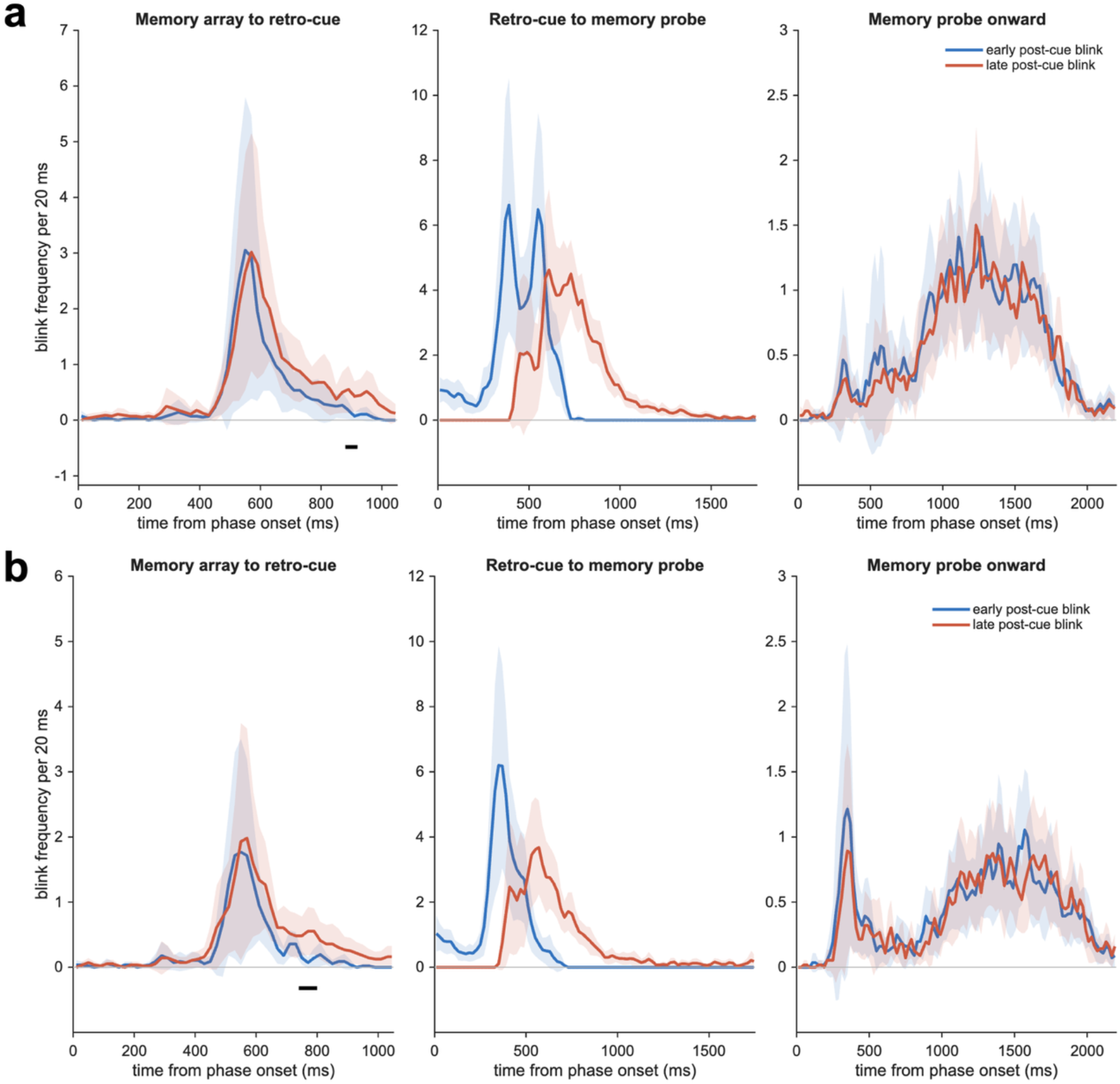
Temporal distribution of first blinks following an early-versus-late post-cue median split. Trials were classified separately within each participant and retro-cue condition according to the latency of the first post-cue blink. **(a)** Selective retro-cue condition. **(b)** Neutral retro-cue condition. Curves show mean blink frequencies in 20-ms bins. Shaded areas indicate 95% confidence intervals around the mean across participants. Black bars denote significant early-versus-late differences from paired two-sided cluster-based permutation tests in the first and third trial phases.

The present analysis addressed an additional, simpler sequential explanation of the post-probe blink frequency pattern. On average, post-cue blinks occurred later in the selective retro-cue condition than in the neutral condition. It was therefore possible that the reduced blink response after the probe was merely a consequence of having blinked more recently before probe onset. Under such an account, trials with a late post-cue blink should show a lower post-probe blink probability, or a delayed post-probe blink distribution, relative to trials with an early post-cue blink. However, the observed post-probe distributions speak against this explanation. The early- and late-blink categories did not produce the pattern expected from a straightforward blink-recency or refractory account, as their temporal profiles did not differ substantially within either retro-cue condition. These results rule out the possibility that the condition difference following the memory probe is simply inherited from the preceding difference in post-cue blink timing between retro-cue conditions.

The analysis nevertheless indicated that post-cue blink classification was not entirely independent of blinking earlier in the trial. The temporal distribution of first blinks following memory array onset differed modestly between trials subsequently classified as containing an early or late post-cue blink. Thus, the timing of blinking after the cue was to some extent related to the preceding blink pattern. This relationship may reflect temporal dependencies or autocorrelation in spontaneous blink behaviour across adjacent phases of a trial. However, it does not compromise the central conclusion of the analysis. Although the post-cue categories were associated with some differences before cue onset, these categories did not produce the inverse relationship between post-cue and post-probe blink distributions required by a refractory explanation.

### 3. Blink latency shuffling approach illustrated by the example of blink-locked event-related potentials in Experiment 2

To illustrate the rationale of the latency-shuffling procedure used for the main blink-locked lateralisation analyses in Experiment 1, we examined blink-locked ERP activity in Experiment 2 at four midline electrodes (Fz, Cz, Pz, POz) before (see Fig. S3 A) and after subtracting the participant-specific shuffled estimate (see Fig. S3 B). Because blinks were not uniformly distributed across the trial, task-related ERP activity can enter the blink-locked signal simply because blinks occur more often during some task periods than others. The shuffled estimate captures this task-related contribution by preserving the participant-specific distribution of blink latencies within the trial while disrupting the trial-specific alignment between blink timing and the EEG signal. Comparing real blink-locked activity with this shuffled estimate therefore tests whether the observed signal exceeds what would be expected from the non-uniform temporal distribution of blinks alone.

**Figure S4.**
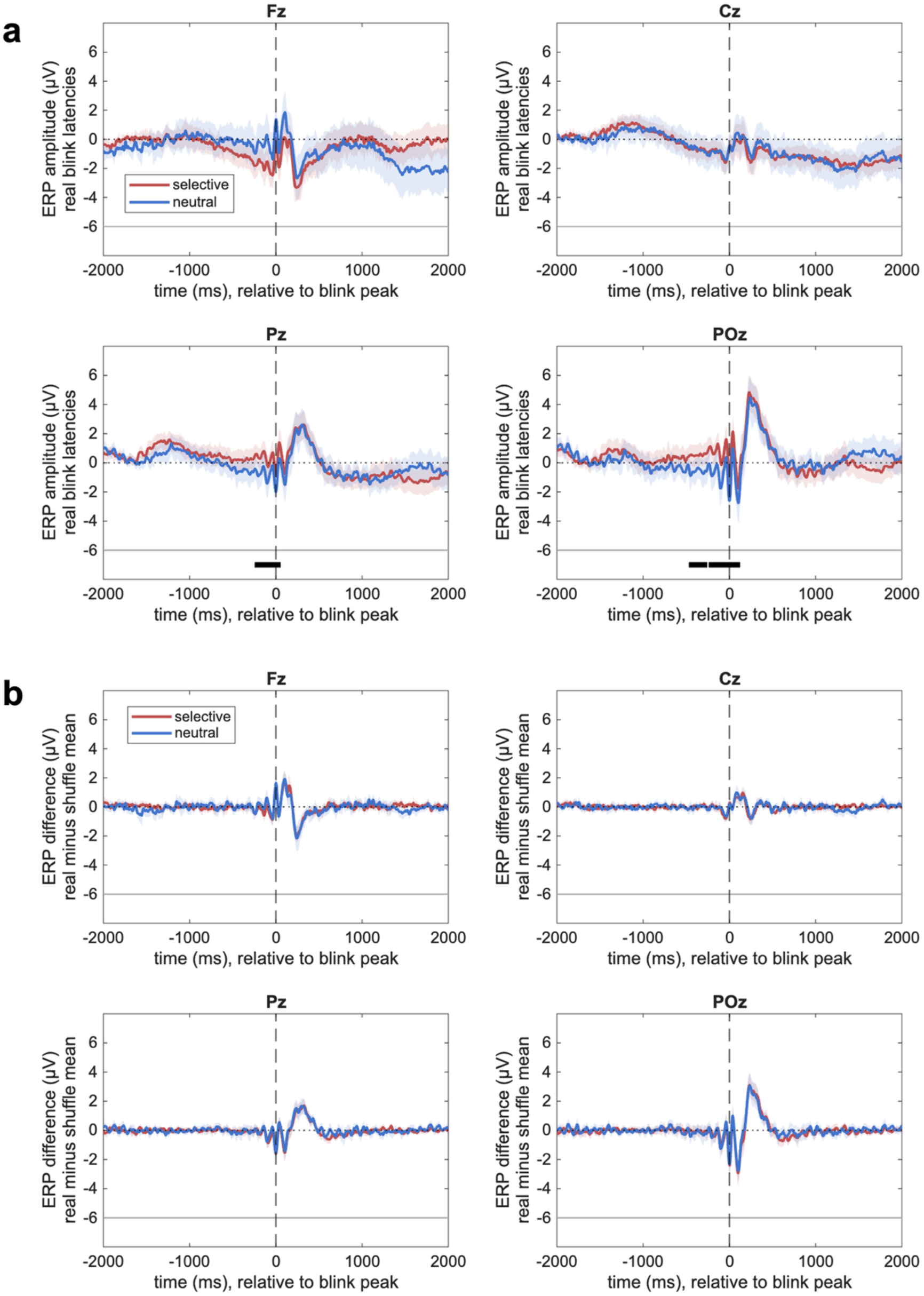
Illustration of the latency-shuffling correction using blink-locked ERPs in Experiment 2. Blink-locked ERP waveforms are shown at four midline electrodes (Fz, Cz, Pz, POz), separately for the selective and neutral retro-cue conditions. **(a)** Real blink-locked ERP waveforms before subtracting the shuffled estimate. Because blinks were non-uniformly distributed across the trial, task-related ERP activity was represented in the blink-locked average. **(b)** Real-minus-shuffled ERP waveforms after subtracting the participant-specific mean latency-shuffled estimate. This subtraction strongly reduced the slow task-related ERP activity and left a temporally focal blink-evoked response centred around the blink peak. Shaded areas indicate 95% confidence intervals around the mean across participants. The dashed vertical line marks the blink peak. Black horizontal bars indicate significant clusters comparing selective and neutral retro-cue conditions.

This logic is very well illustrated in the ERP waveforms. Before subtracting the shuffled estimate, blink-locked ERPs showed slow task-related activity that differed between the selective and neutral retro-cue conditions, particularly at posterior electrodes. Such condition differences are expected because selective and neutral retro-cues are known to evoke different slow ERPs during working memory maintenance ^1^. Importantly, these task-related differences were strongly reduced after subtracting the shuffled estimate. The remaining real-minus-shuffled waveforms were dominated by a temporally focal blink-evoked response centred around the blink peak and did not show statistically significant differences between the selective and neutral retro-cue conditions.

Thus, this supplementary analysis illustrates what the shuffled control achieves in the main blink-locked analyses. It removes EEG activity that is carried into the blink-locked average by the non-uniform timing of blinks within the task, including known cue-related slow-wave ERP differences. Consequently, when real and shuffled blink-locked lateralisations are compared in the main analyses (see Fig. 2 in the manuscript), the resulting effects cannot be explained simply by the fact that blinks occur preferentially during task periods with strong task-related EEG activity. Rather, they indicate signal components that are more specifically aligned with the true timing of blinks relative to the participant-specific stimulus-locked expectation.

